# Distinct filament morphology and membrane tethering features of the dual FtsZs in Odinarchaeota

**DOI:** 10.1101/2025.02.03.636245

**Authors:** Jayanti Kumari, Akhilesh Uthaman, Ananya Kundu, Anubhav Dhar, Vaibhav Sharma, Sucharita Bose, Soumyajit Dutta, Srijita Roy, Ramanujam Srinivasan, Samay Pande, Kutti R. Vinothkumar, Pananghat Gayathri, Saravanan Palani

## Abstract

The deep-branching Asgard archaea have emerged as a model to study Eukaryogenesis as they harbor homologs of many of the major eukaryotic protein families. In this study, we use Candidatus Odinarchaeota FtsZs as representatives to understand the probable origin, evolution, and assembly of the FtsZ-Tubulin protein superfamily in Asgard archaea. We have performed a comparative analysis of the biochemical properties and cytoskeletal assembly of FtsZ1 and FtsZ2, the two FtsZ isoforms in the Odinarchaeota metagenome. Our electron microscopy analysis reveals that OdinFtsZ1 assembles into curved single protofilaments, while OdinFtsZ2 forms stacked spiral ring-like structures. Upon sequence analysis, we identified an N-terminal amphipathic helix in OdinFtsZ1, which mediates direct membrane tethering. In contrast, OdinFtsZ2 is recruited to the membrane by the membrane-anchor OdinSepF via OdinFtsZ2’s C-terminal tail. Overall, we report the presence of two distant evolutionary paralogs of FtsZ in Odinarchaeota, with distinct filament assemblies and differing modes of membrane targeting mechanisms, namely direct vs SepF-mediated membrane binding. Our findings highlight the diversity of FtsZ proteins in the transitional archaeal phylum Odinarchaeota, providing valuable insights into the evolution and differentiation of tubulin family proteins.

## Introduction

Asgard archaea, a newly identified archaeal superphylum, has emerged as the closest relative of Eukaryotes^1–4^. Metagenome Assembled Genome (MAGs) of these archaea revealed the presence of multiple Eukaryotic Signature Proteins (ESPs) such as cell-division-associated proteins, trafficking proteins, protein translocation and glycosylation-related proteins^2,5^. The identification of cytoskeletal protein homologs like Lokiactin, gelsolin, profilin, and homologues of subunit 4 of the Arp2/3 complex in various Asgard lineages provides evidence for the archaeal origin of the present day eukaryotic cytoskeleton^6–10^.

An interesting feature of Asgard archaea is the presence of both prokaryotic (FtsZ, SepF) and eukaryotic (tubulin, ESCRT, actin) cytoskeletal elements, raising questions on their organization and role in processes such as cell division. FtsZ, a tubulin superfamily protein, is an essential player in cell division in many bacteria and archaea^11–14^. In bacteria, overlapping FtsZ filaments assemble as Z-rings at the cell midbody and help recruit other divisome proteins^13^. Tread-milling filaments of FtsZ form the leading edge of the invaginating septum and direct inward growth of the septal wall to ensure the successful separation of cells^12–16^. FtsZ filament formation is GTP-dependent, and results in a GTP-hydrolysis site at the interface of two monomers^20–24^, as is the case with α/β-tubulin^25,26^. However, the polarity of FtsZ filaments is expected to be opposite to that of tubulin, with subunits being added at the bottom end (defined as the exposed interface consisting of C-terminal Domain (CTD)), away from the nucleotide-binding pocket^24,26^. Several studies, including electron cryotomography of *Caulobacter crescentus* cells, revealed arc-like FtsZ filaments at mid-cell^15^. Additionally, investigation of in vitro reconstituted FtsZ filaments inside liposomes shows the presence of ring-like structures, which can cause membrane deformation upon GTP hydrolysis^27–29^ This ring is anchored to the membrane via another divisome protein called FtsA, an actin homologue^30–32^. In addition to FtsA, many Gram-negative bacteria, including *Escherichia coli*, possess additional FtsZ membrane anchors like ZipA^33^, whereas Gram-positive bacteria and cyanobacteria possess a different anchor called SepF. SepF has been reported to bind FtsZ, and interestingly is conserved in archaeal systems as well^34–37^.

Unlike most bacteria which depend solely on FtsZ for cell division^11-13^, archaeal superphylum harbors both FtsZ^14^ and ESCRT homologs^38–41^. Archaeal cell division, outside of Asgard superphylum, depends on either one or two forms of FtsZ, with SepF acting as a membrane anchor^36^. The archaeal cell division mechanism and the key factors involved remain highly unexplored. Asgard archaeon Odinarchaeota uniquely possesses both a tubulin-like gene and two FtsZ genes^42,43^. A recent structural study on OdinTubulin and comparison to the eukaryotic α/β-tubulins has suggested the possible evolution of tubule-like structures via stabilization of the inter-protofilament interactions^43^. This intriguing co-existence of ancestral FtsZ and tubule-forming tubulin, among Asgard archaea (that are closest relatives to eukaryotes), raises interesting questions about the origins of eukaryotic tubulin, their adaptation into chromosome segregation, cytoskeletal arrangement and function in these archaea, and their respective contributions to the cell division process. The study of Odin cytoskeletal proteins, thus, holds the potential for unveiling new insights into crosstalk and coordination among cytoskeletal protein families of Asgard archaea and their gradual evolution that led to the present eukaryotic cytoskeleton. Although the structure of OdinTubulin is known, the structural features and functional specializations of the two FtsZ proteins in Odinarchaeota remain completely unknown despite their existence alongside other cytoskeletal elements in a single organism. Moreover, diversification of functions has also occurred during evolution among the FtsZ/tubulin family of proteins. For example, TubZ^44^ and PhuZ^45,46^ are involved in DNA segregation and maintenance in bacteria and bacteriophages, respectively. On the contrary, The CetZ family of proteins that are commonly found in many archaea play a role in cell shape maintenance and do not affect cell division processes^47^. Further, MinD, a spatial regulator of Z-ring assembly in *E. coli*, has been co-opted in archaea to position the cell motility and chemosensory receptors^48^. These led us to carry out studies on the two candidate FtsZ genes in Odin to characterize their biochemical properties and filament structures, and to gain insights about their functions.

Here, we present evidence that two FtsZ-like genes in Candidatus Odinarchaeota assemble into distinct filament morphologies in the presence of GTP. OdinFtsZ1 assembles into typical FtsZ single protofilament structure whereas OdinFtsZ2 assembles as spiral filament morphology. We show that SepF, a characteristic membrane anchor for FtsZs, can specifically bind OdinFtsZ2 via the C-terminal variable region, whereas OdinFtsZ1 associates with membranes via its hydrophobic N-terminal region that enables direct membrane binding independent of SepF. Overall, our study provides insights into the biochemical and structural features of dual FtsZs of the Asgard archaea and their potential roles as part of the cell division apparatus in these closest relatives of eukaryotes, with implications for the origin/diversification of the Tubulin superfamily of proteins across the kingdoms of life.

## Results

### Candidatus Odinarchaeota encodes two FtsZ homologs

A phylogenetic analysis of FtsZ protein sequences from the Asgard group identified two candidate sister clades corresponding to two distinct clusters in Odinarchaeota representing FtsZ1 and FtsZ2 proteins. This distribution indicates independent evolutionary trajectories (Fig. S1A). AlphaFold^49^-predicted monomeric structures of OdinFtsZs (UNIPROT Accession: OdinFtsZ1 (A0AAF0ID74), OdinFtsZ2 (A0AAF0D3V8)) show that both the candidate proteins, like bacterial FtsZ (*E*.*coli* FtsZ structure shown; PDB: 6UMK) and tubulin, possess an N-terminal domain, which consists of the nucleotide-binding pocket and a C-terminal domain with the H7 helix positioned between the two domains (Fig. 1A). The end of this helix carries a loop corresponding to the catalytic loop important for GTP hydrolysis in bacterial FtsZ proteins (Fig. 1A). The presence of highly conserved GTP binding (GGGTG[T/S]G) and GTP hydrolysis (NxDxx[D/E]) motifs^50^ in these archaeal sequences indicates conservation of crucial functional elements across the tree of life (Fig. 1B). Though OdinFtsZ2 clusters along with other FtsZ2 members (Fig. S1A), it possesses an insertion of a loop in the CTD, which is not found in other FtsZ proteins (Fig. 1A, S1B).

**Figure 1:**
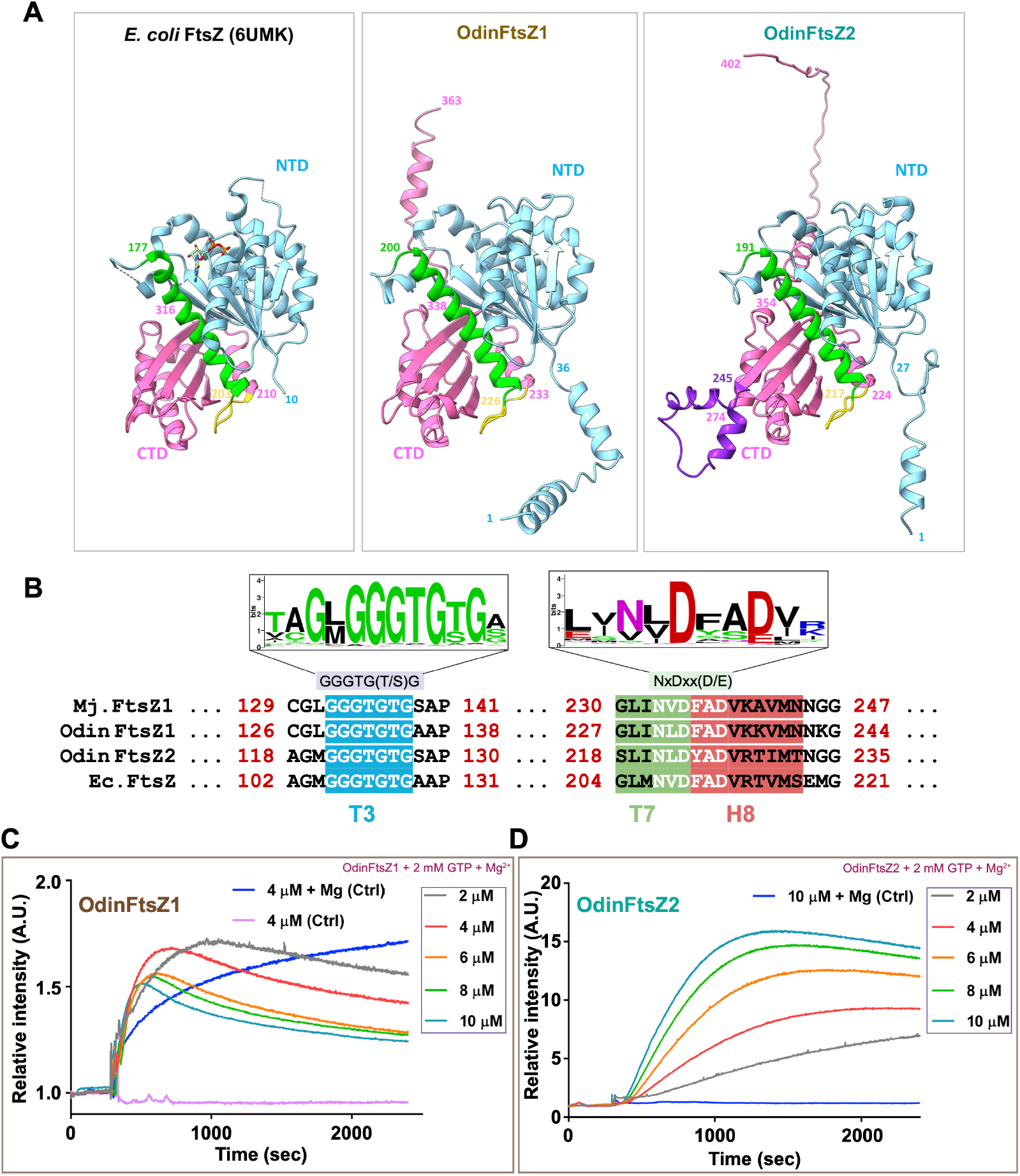
Odin from Asgard archaea carries two paralogs of polymeric FtsZ proteins. **(A)** Structural comparison of protomers predicted by AlphaFold for OdinFtsZ proteins with EcFtsZ monomeric structure (PDB-6UMK). Cyan represents the N-terminal domain of the structure, green denotes the H7 helix, yellow denotes the T7 loop, pink denotes the C-terminal domain of the FtsZ monomer, and purple is the extra loop insertion in OdinFtsZ2. **(B)** Representative sequence logo^14^ for the tubulin signature motifs: GTP binding (left) and GTP hydrolysis motif (right). **(C)** The plot of relative light scattering intensity (y-axis) with time (x-axis) for different concentrations of OdinFtsZ1 protein polymerized in the presence of 2 mM GTP and 5 mM Mg^2+.^ The relative scattering intensity of the protein at 4 μM with Mg^2+^ (in blue) and without Mg^2+^ (in magenta) but devoid of any nucleotide act as controls. **(D)** The plot of relative light scattering intensity (y-axis) with time (x-axis) for OdinFtsZ2 protein polymerized in the presence of 2 mM GTP and 5 mM Mg^2+^. The blue curve represents 10 μM protein polymerized solely in the presence of 5 mM Mg^2+^, which serves as the control.

FtsZ, being an ancestral relative of tubulin, polymerizes in a GTP-dependent manner^51–53.^ To monitor the polymerization of purified OdinFtsZ1 and OdinFtsZ2 proteins (Fig. S1C), we employed 90^°^ angle light scattering^54^. For the OdinFtsZ proteins, there was an increase in scattering upon the addition of GTP. Interestingly, the maximum relative intensity of polymerized OdinFtsZ1 exhibited a marginal decrease, while the maximum intensity was achieved faster, with increasing protein concentration (Fig. 1C). This observation could possibly be attributed to an enhanced number of nucleation events at higher protein concentrations. Increased nucleation events likely lead to the formation of numerous short filaments, thus limiting the overall relative intensity. Notably, there was an increase in scatter observed in OdinFtsZ1 in the presence of Mg^2+^ ion (5 mM), even without any supplemented nucleotide, a property not observed for OdinFtsZ2. In the presence of saturating GTP levels, OdinFtsZ2 showed an apparent increase in the maximum relative intensity with increasing protein concentration (Fig. 1D). The observed variation in light scattering associated with FtsZ polymerization might be linked to polymer mass^54^ or to distinct filament morphologies, and assembly dynamics for FtsZ1 and FtsZ2.

### OdinFtsZ1 and OdinFtsZ2 interact despite distinct filament assembly

The difference in the scattering pattern between the two proteins suggests potential disparities in filament assembly. Therefore, we employed cryogenic-sample electron microscopy (Cryo-EM) to visualize and compare the filament morphology. We assessed the ability of the protein to form filaments in the presence of GTP. We observed filaments for OdinFtsZ1 (Fig. 2A, B), similar to the single protofilaments of other canonical FtsZs^55–58^. To understand how different nucleotide states affect the polymerization of these filaments, we tested the assembly of OdinFtsZ filaments in the presence of different nucleotide analogs using negative staining and transmission electron microscopy (TEM). We observed that OdinFtsZ1 forms short filaments in the presence of GTP and-non-hydrolysable GTPγS and GMPPCP (a slow hydrolyzing nucleotide analogue) (Fig. S2A). OdinFtsZ1, even without added nucleotide, formed short filaments (Fig. S2A).

**Figure 2:**
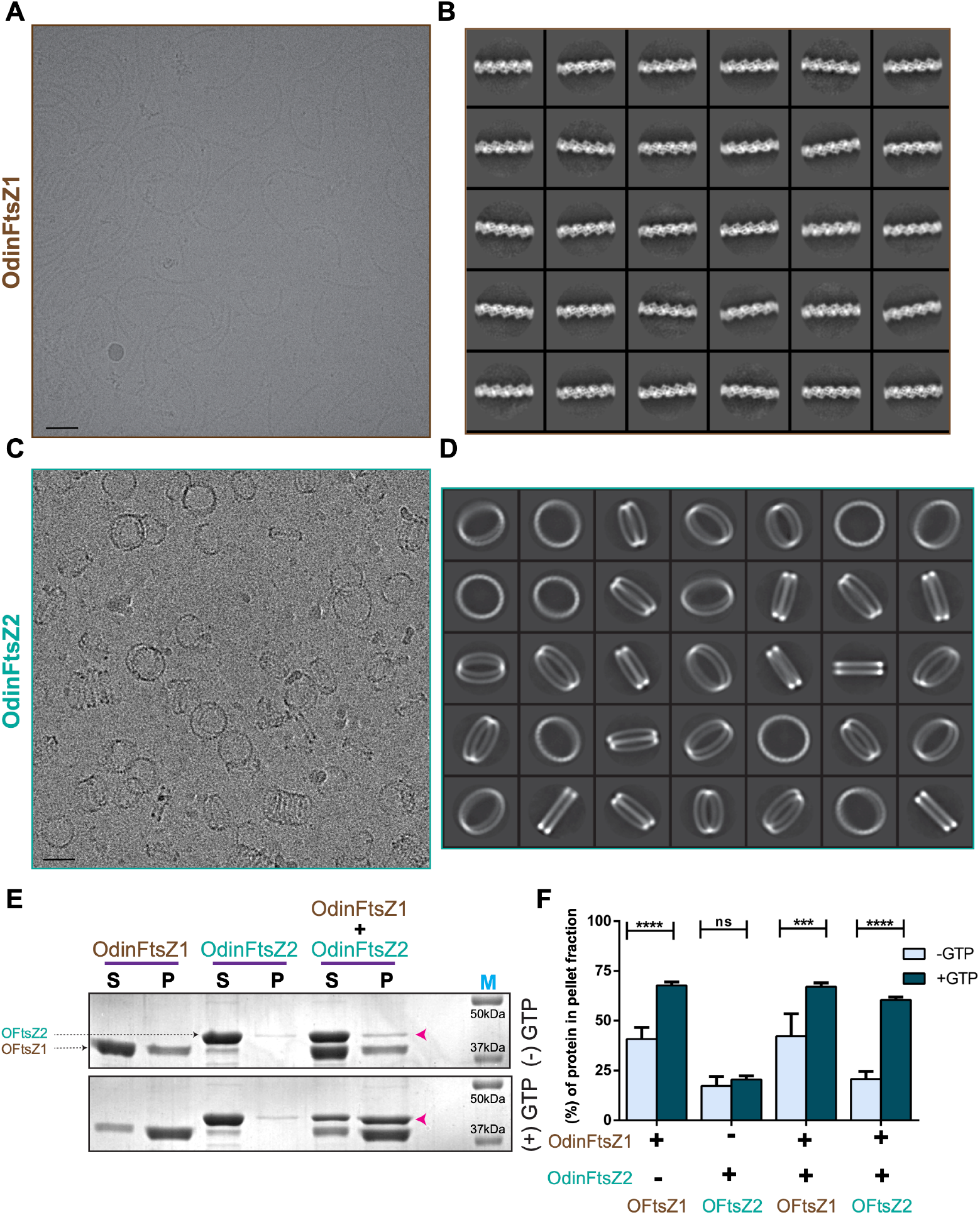
OdinFtsZ proteins assemble into distinct filament morphology and co-pellets on ultracentrifugation. **(A)** Cryo micrograph for OdinFtsZ1 (10 μM) polymerized in the presence of 2 mM GMPCPP (scale bar – 500 Å). **(B)** Representative 2D average class for OdinFtsZ1. **(C)** Cryo micrograph for OdinFtsZ2 polymerized with 2 mM GTP (scale bar – 500 Å). **(D)** Representative 2D class averages for particles selected only for shorter stacks. **(E)** Representative 12% SDS-PAGE gel for sedimentation of individual proteins OdinFtsZ1 (10 μM), OdinFtsZ2 (10 μM) and the co-sedimentation when incubated in equimolar concentration (10 μM) in the absence (top) and the presence of GTP nucleotide (bottom). Both supernatant (S) and pellet (P) fractions upon ultracentrifugation at 100,000 ×*g* were loaded. **(F)** Representative plot for the mean percentage of protein in the pellet fraction (y-axis) with and without GTP. The pellet (or supernatant) fraction intensities corresponding to the OdinFtsZ1 band were calculated as the intensity of the pellet (or supernatant) divided by the sum of the two intensities and represented in the graph as a percentage. ‘+’ or ‘-’ below the bars denote the presence/absence of OdinFtsZs, while the OFtsZ1/OFtsZ2 labelled below the bottom lines denote the protein band which has been quantified. (N=3, Two-Way ANOVA with Sidak’s multiple comparisons test was used, ^ns^ non-significant, *** p -0.002, **** p < 0.0001).

Interestingly, Cryo-EM of OdinFtsZ2 protein revealed that the filaments self-assembled into structures with high curvature resembling stacked spring-like assemblies (Fig. 2C). OdinFtsZ2 formed highly curved, ring-like structures when provided with GTP or GTPγS while showing very little to no filament assembly in the presence of GMPCPP and GDP (Fig. S2B). No rings were observed when OdinFtsZ2 was devoid of any excess nucleotide, which suggests the role of nucleotide binding for the assembly (Fig. S2B). This observation is consistent with the absence of an increase in light scattering when there was no nucleotide addition (Fig. 1D). A key observation is the formation of a variable number of rings in the stacks by OdinFtsZ2 in the current experimental conditions (Fig. 2C), similar to the OdinTubulin^43^. In a given field of view, the number of longer stacked rings was less, but shorter stacks were more prominent. These were picked and averaged in 2D (Fig. 2D). Unlike the 2D class averages of OdinFtsZ1, which show high-resolution features, the 2D classes of OdinFtsZ2 show low-resolution features of the ring and no clear indication of subunit stoichiometry indicating heterogeneity (Fig. 2B and 2D respectively). Factors that promote a stable formation of stacked rings are currently being investigated.

To assess if OdinFtsZ proteins interacted with each other and formed co-polymers, we utilized polymer pelleting or sedimentation assays. FtsZ polymerization and subsequent pelleting were first tested individually for OdinFtsZ1 and OdinFtsZ2 both in the presence and absence of GTP. Again, consistent with the light-scattering and TEM micrographs, we observed a substantial pellet fraction for OdinFtsZ1 without supplementing any nucleotide (Fig. 2E). However, adding GTP significantly enhanced the amount of OdinFtsZ1 in the pellet fraction, suggestive of a filament assembly stimulated in the presence of nucleotide. However, OdinFtsZ2 was not observed in the pellet fraction, irrespective of the presence or absence of GTP (Fig. 2E). It is interesting to note that OdinFtsZ2 does not sediment despite the formation of higher-order assemblies confirmed by TEM and scattering.

Since OdinFtsZ2 failed to pellet when incubated with GTP, we could use this assay to test the probable co-pelleting of OdinFtsZ2 in the presence of OdinFtsZ1. When both the OdinFtsZs were incubated at equimolar concentrations in the presence of GTP, a significant fraction of OdinFtsZ2 was seen in the pellet fraction along with OdinFtsZ1 (Fig. 2E, 2F). This co-sedimentation was strictly dependent on the presence of nucleotides, suggestive of an interaction between the two FtsZ homologs, probably between the filament states. This interaction between OdinFtsZ1 and OdinFtsZ2 observed in vitro suggests a potential interaction in vivo as well as a coordinated cellular function for the FtsZs in Odinarchaeota.

### SepF mediated membrane tethering of OdinFtsZ2

In bacteria, FtsZ proteins are typically tethered to the membranes via interactions through their conserved CCTP motifs with adaptor proteins like FtsA, ZipA and SepF ^33,36,59^. While ZipA is an integral membrane protein^33^, the anchor proteins SepF and FtsA (an actin homolog) bind the membrane via amphipathic helices at their N-termini and C-termini, respectively^36,59^. In contrast to FtsA, SepF is widely conserved in the archaeal superphylum (Fig. S3A). The consistent presence of SepF in archaeal systems possessing FtsZ implies a strong functional interdependence between these proteins. Recent studies report that these proteins localize to the Z-ring and that their depletion causes severe cell division defects^37,34^. SepF dimers from bacterial systems like *B. subtilis* are known to oligomerize further^60^. Residues important for these interactions are not conserved in other bacterial species such as *C. glutamicum*^61^ and archaeal SepF^36^. *In-vitro* studies support no further oligomeric assembly for archaeal SepF and thus their oligomerization status remains to be tested. To validate the conserved role of SepF in Asgard, we investigated the membrane-binding ability of OdinSepF (UNIPROT Accession: A0AAF0D343). The helical wheel diagram represents a canonical N-terminal amphipathic helix in OdinSepF protein (Fig. S3B). We assessed its membrane interaction via liposome-based pelleting. OdinSepF exhibited strong binding to liposomes as observed by a considerable amount of SepF in the pellet fractions with increasing liposome concentration during ultracentrifugation (Fig. S3C, S3D). The oligomeric status of OdinSepF was tested by size exclusion chromatography with multi-angle light scattering (SEC-MALS). This enabled the accurate mass measurement of OdinSepF corresponding to the molecular weight of the dimeric protein (Fig. S3E) as observed for other archaeal SepF proteins^36^.

To check for the interaction between OdinSepF and OdinFtsZ1/Z2, we employed a pull-down assay using purified proteins (Fig. S3F). Our result indicated interaction between OdinFtsZ2 and SepF, while no detectable interaction was observed with OdinFtsZ1 (Fig. 3A). In most archaeal species, this binding is mediated by a nearly conserved GID motif present at the C-terminus of FtsZ^36^. Unlike OdinFtsZ1, OdinFtsZ2 contains an extra C-terminal stretch, which might possess a motif analogous to the GID motif (Fig. S3G). To further confirm the role of the C-terminus, we constructed a truncated OdinFtsZ2 lacking the C-terminal region (OdinFtsZ2^ΔC49^). This construct completely abolished OdinSepF binding, suggesting a specific interaction between OdinSepF and OdinFtsZ2 through the FtsZ2 C-terminus (Fig. 3B). To validate the role of this interaction for OdinFtsZ2 membrane anchoring, we carried out co-sedimentation assays of SepF-OdinFtsZ2 mixtures in the presence of liposomes. OdinFtsZ2, which exhibits no independent binding with liposomes, co-pelleted with liposomes when incubated with its membrane anchor, SepF (Fig. 3C, 3E). This suggests that OdinFtsZ2 can be indirectly tethered to membranes through its interaction and association with OdinSepF. The deletion of C-terminal, OdinFtsZ2^ΔC49^, abolished its interaction with SepF and the protein remained in the supernatant regardless of the presence of liposomes (Fig. 3D, 3F).

**Figure 3:**
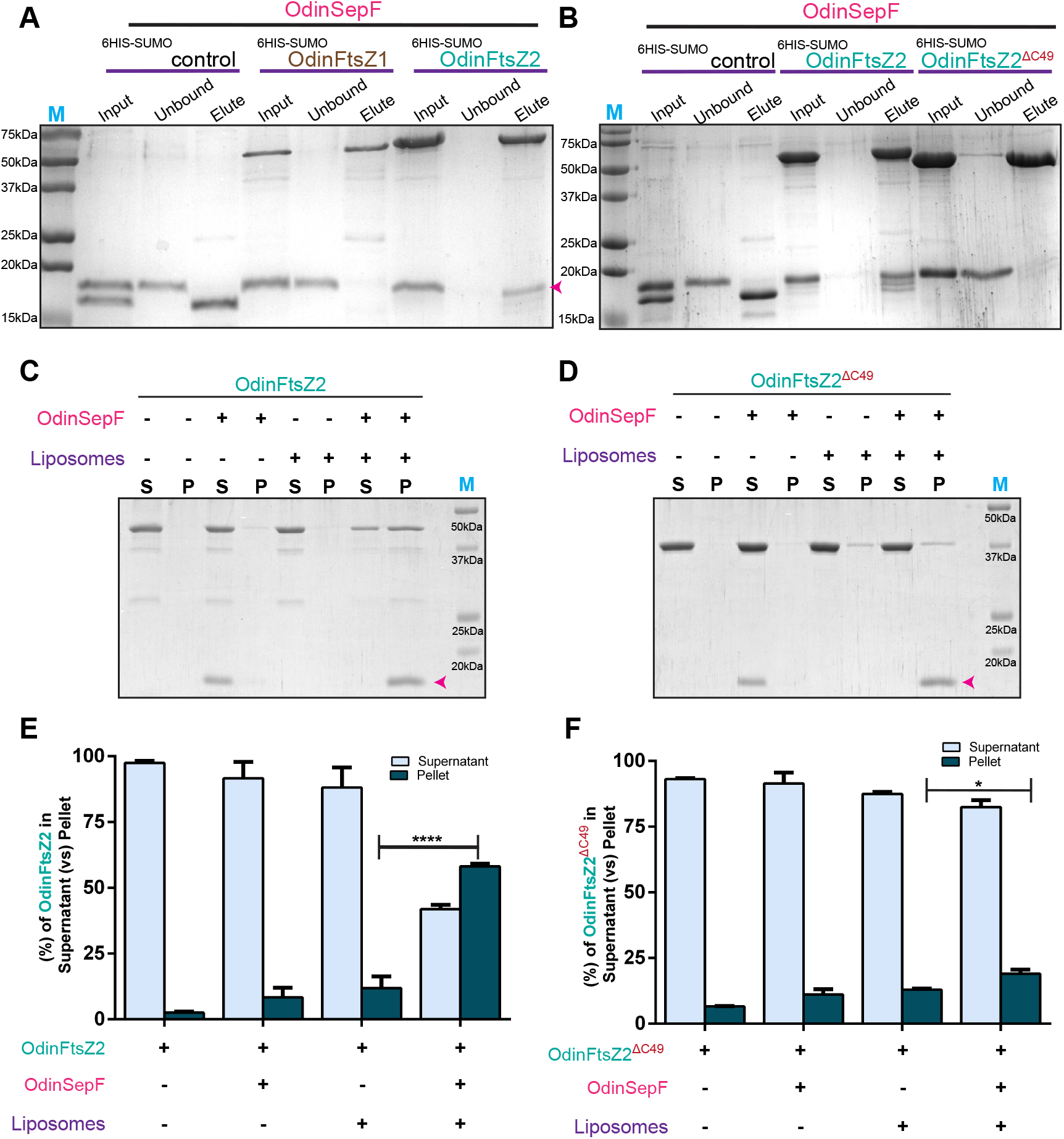
OdinFtsZ2 is recruited to liposome via OdinSepF, whereas OdinFtsZ1 shows no physical interaction with OdinSepF. **(A)** Representative 15% SDS-PAGE gel of in-vitro binding for 6HIS-SUMO, OdinFtsZ1 and OdinFtsZ2. Bead-bound 6HIS-SUMO, OdinFtsZ1 and OdinFtsZ2 were incubated with untagged OdinSepF. Input, wash and elution fractions were loaded after the incubation. The pink arrow denotes the presence of OdinSepF in the elution fraction of OdinFtsZ2. **(B)** Representative 15% SDS-PAGE gel of in-vitro binding for 6HIS-SUMO, OdinFtsZ2 and OdinFtsZ2^ΔC49^. **(C)** Representative 12% SDS-PAGE gel for liposome co-sedimentation of OdinFtsZ2 in the presence of OdinSepF. The supernatant (S) and pellet (P) fractions upon sedimentation at 100,000 ×*g* were analyzed for OdinFtsZ2 (alone), OdinFtsZ2 + OdinSepF, OdinFtsZ2 + Liposomes, OdinFtsZ2 + OdinSepF+ Liposomes. The arrow represents the presence of OdinSepF in the respective lanes. **(D)** Representative 12% SDS-PAGE gel for liposome co-sedimentation of OdinFtsZ2^ΔC49^ in the presence of OdinSepF. The supernatant (S) and pellet (P) fractions upon sedimentation at 100,000 ×*g* were analyzed for OdinFtsZ2^ΔC49^ (alone), OdinFtsZ2^ΔC49^ + OdinSepF, OdinFtsZ2^ΔC49^ + Liposomes, OdinFtsZ2^ΔC49^ + OdinSepF + Liposomes. **(E)** Plot representing the percentage fraction of OdinFtsZ2 (y-axis) in the supernatant (S) and pellet (P) fractions (Fig. 3C). (N=3, Two-Way ANOVA with Tukey’s multiple comparisons test was used, ^ns^ non-significant, **** p < 0.0001) **(F)** Plot representing the percentage fraction of OdinFtsZ2^ΔC49^ (y-axis) in the supernatant (S) and pellet (P) fraction (Fig.3D). (N=3, Two-Way ANOVA with Tukey’s multiple comparisons test was used, ^ns^ non-significant, * p - 0.0309).

### Direct membrane anchoring via an N-terminal amphipathic helix in Odin FtsZ1

Reports on *H*.*volcanii* revealed that FtsZ1 is one of the earliest proteins to localize at the future division site. It is hypothesized that FtsZ1 recruits other divisome proteins to the site of cell division, which in turn organizes SepF^37^. Pull-down assays in these organisms as well as in Odin, support no physical interaction of SepF with FtsZ1. The mechanism by which FtsZ1 assembles at the future division site remains unidentified. Recent studies on Mycoplasma FtsZs have reported the presence of putative membrane targeting sequences (MTS), which are capable of binding to the membrane^62^. Also, chloroplast FtsZs are known to interact directly with the membrane^63,64^. To investigate the possibility of a similar mechanism we analyzed the sequence of OdinFtsZ proteins. This revealed the presence of an N-terminal extension unique to OdinFtsZ1 that was absent in OdinFtsZ2. The representative helical wheel diagram depicts a potential amphipathic helix at the N-terminal of OdinFtsZ1(Fig. 4A). We hypothesized that this helix might possess membrane binding ability. This membrane interaction was tested with liposome pelleting assay where OdinFtsZ1 showed a significant amount in the pellet fraction in the presence of liposomes. (Fig. 4B-D). An N-terminal deletion construct (OdinFtsZ1-ΔNTAH) lacking the first 14 amino acids abolished the membrane binding, suggesting the presence of an amphipathic helix at the N-terminus which helps in mediating direct membrane attachment (Fig. 4C).

**Figure 4:**
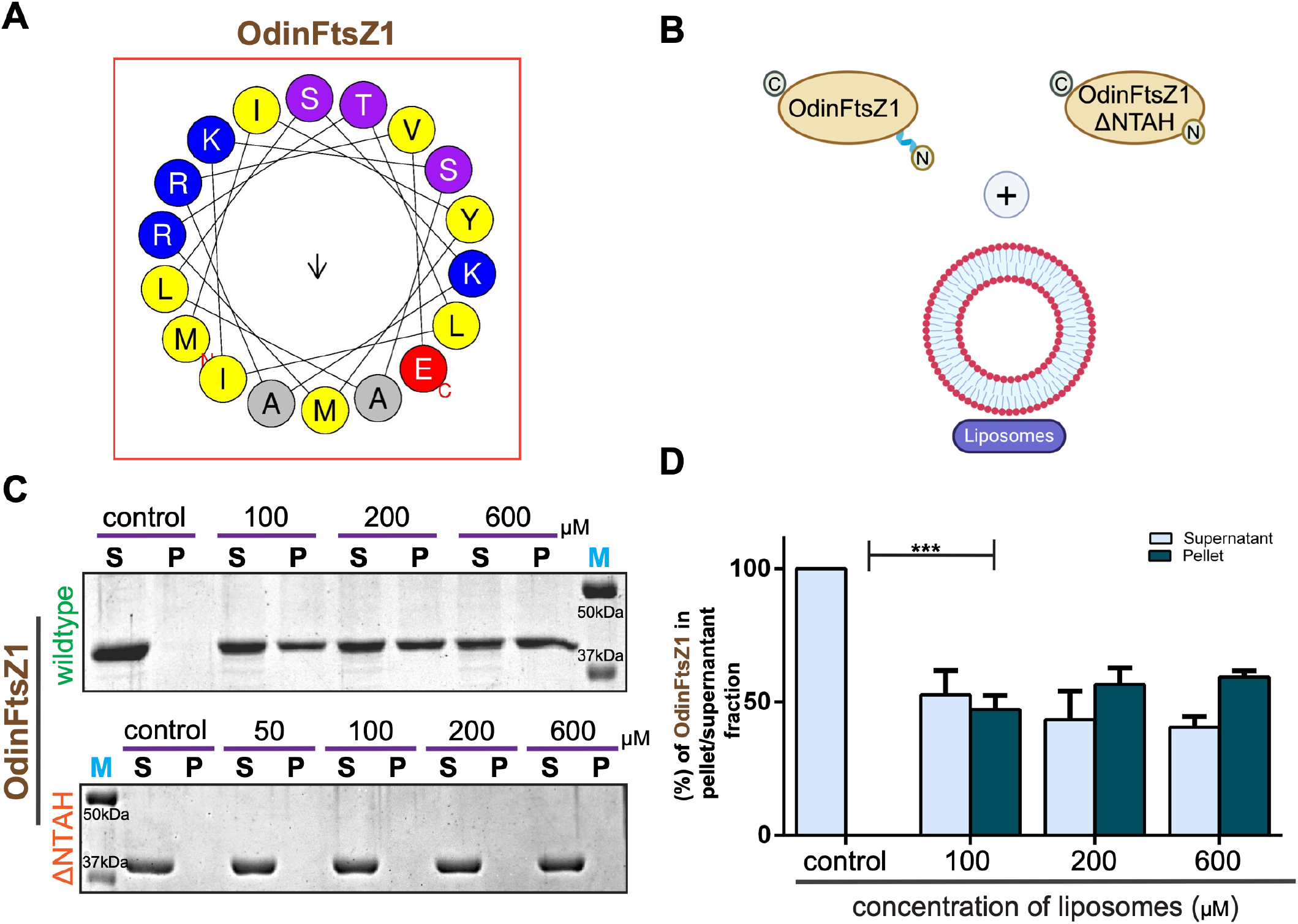
OdinFtsZ1 binds liposomes via its N-terminal extension. **(A)** Representative helical wheel diagram for the predicted amphipathic helix (AH) at the N-terminus of OdinFtsZ1 screened using Heliquest (version 1.2 Analysis module)^68^. **(B)** Schematic representation for liposome binding by OdinFtsZ1. **(C)** Representative 12% SDS-PAGE gel for the sedimentation of OdinFtsZ1 with increasing concentration of liposome. The supernatant (S) and pellet (P) fractions were analyzed and loaded upon sedimentation at 100,000 ×*g*. Lanes 1 and 2 represent S and P for only OdinFtsZ1 without liposomes. **(D)** Representative plot for the percentage fraction of OdinFtsZ1(y-axis) in the supernatant (S) and pellet (P) fractions with increasing concentration of liposomes as represented in Fig. 4C. (N=3, Two-Way ANOVA with Tukey’s multiple comparisons test was used, ^ns^ non-significant, *** p-0.0001)

## Discussion

FtsZ is a key protein essential for cell division in bacterial ^11,12^ and many archaeal systems^14^. Over evolutionary time, these proteins have transitioned into actin/tubulin-like proteins during eukaryogenesis and subsequent divergence^65^. It can be speculated that this evolutionary shift within archaea has led to a transition towards an ESCRT-based cell division mechanism with a loss of FtsZ genes in several clades. Interestingly, evidence of this shift is indicated by the presence of both FtsZ and tubulin-like proteins in certain clades (e.g. Odinarchaeota) of the Asgard superphylum. This unique dual existence of these proteins within Odinarchaeota offers an evolutionary window to study and understand the gradual transition from FtsZ to tubulin-like proteins, leading to the diversification of functions in the FtsZ/tubulin superfamily.

Our study aimed to investigate the distinct structural and biochemical features of the dual FtsZs from Odinarchaeota. Our findings reveal the presence of two distinct FtsZ proteins in Odinarchaea, each exhibiting unique filament assembly characteristics. OdinFtsZ1 showed canonical protofilaments as reported for many bacterial FtsZs, suggesting a conserved mode of assembly across kingdoms. On the other hand, OdinFtsZ2 assembled into stacked-ring-like structures which have a striking resemblance to filament morphologies adopted by OdinTubulin^43^. Since OdinFtsZ2 and OdinTubulin share a low similarity in terms of sequence and structure, similar filament morphologies may indicate their involvement in a common process or function. The observed difference in the filament morphologies of the two FtsZ proteins suggests probable functional specialization within archaeal cells, while their in-vitro interaction hints at a potential collaborative role in cellular processes. We also note the similarity of the OdinFtsZ2 rings to the recently reported FtsA rings that prevent FtsZ filaments from bundling and the cryo-EM structures of FtsZ-FtsA filaments reconstituted within liposomes^32,66^. This similarity may hint at a similar pairing between OdinFtsZ1 and OdinFtsZ2, which remains to be explored.

Our study also addresses the distinct membrane tethering mechanisms adopted by the dual Odin FtsZs. Interestingly, OdinFtsZ1 possesses an N-terminal amphipathic helix capable of membrane binding like one found in mycoplasma and chloroplast FtsZ. OdinFtsZ2, however, does not possess direct membrane binding but can indirectly bind via its membrane anchor, SepF. SepF is highly conserved among archaea as a membrane anchor for FtsZ and co-exists in all archaea with FtsZ genes^36^. It is interesting to observe that, unlike the bacterial homologues, the archaeal SepF does not form higher-order assemblies, however, it interacts with OdinFtsZ2, which has a stacked ring-like assembly. OdinFtsZ2-SepF interaction is mediated by the C-terminal domain of OdinFtsZ2, a feature that seems to be present across different domains of life^59,64^.

Together, though it is not possible to assign the proteins studied in this study as the direct ancestors of the evolved cytoskeletal elements, our findings support the hypothesis that the eukaryotic cytoskeleton evolved through functional diversification and innovation within the transitional archaeal phylum Odinarchaeota. Importantly, based on our findings, at least four predictions can be made about the evolution of cytoskeletal elements. For example, the distinct roles of OdinFtsZ1 (direct membrane tethering via an N-terminal amphipathic helix) and OdinFtsZ2 (SepF-mediated recruitment), as demonstrated here, reflect an early tendency toward functional specialization within the cytoskeleton. In eukaryotes, this may have laid the groundwork for the evolution of diverse cytoskeletal proteins like tubulin (forming microtubules) and actin, each with distinct roles in cell shape, division, and intracellular transport. Second, the demonstration of the recruitment of OdinFtsZ2 by OdinSepF highlights the early role of accessory proteins in cytoskeletal function, a feature that is extensively elaborated in eukaryotes. Such accessory proteins may have facilitated the evolution of regulatory networks governing cytoskeletal dynamics, which are critical for eukaryotic cellular complexity. Third, dual mechanisms of membrane recruitment in Odinarchaeota (direct tethering via amphipathic helices vs. accessory proteins like SepF) could represent an ancestral strategy that predated the evolution of eukaryotic cytoskeletal-membrane interactions. These mechanisms likely evolved further in eukaryotes to support processes such as endocytosis, intracellular trafficking, and organelle positioning. Finally, our findings suggest that the Asgard archaea, particularly Odinarchaeota, might represent an intermediate stage of cytoskeletal evolution, bridging simple prokaryotic systems (e.g., bacterial FtsZ) and the highly complex eukaryotic cytoskeleton. This intermediate complexity could have provided a scaffold for the emergence of more dynamic and adaptable cytoskeletal elements required for larger, compartmentalized cells.

Future studies should reveal more insights into the cellular functions of the distinct FtsZs in Odinarchaeota. It is possible that like in the archaeon *Haloferax*^14^, the two Odin FtsZs with distinct properties might cooperate in cell division. Interestingly, the two Odin FtsZ genes are located along with protein biosynthesis genes such as the ribosomal proteins and tRNA synthetases^42^ leading to the possibility of co-ordination of cellular biosynthesis with cell division cycles, often observed across species. Alternately, we imagine that the two FtsZs might have distinct functions. OdinFtsZ1 filaments may adopt a classical Z-ring architecture for cell division and OdinFtsZ2 (like CetZ) may play a role in functions such as membrane deformation forming of intra-membrane structures or cell shape maintenance, due to the fact that it shares an extruding helical loop in the C-terminal domain, similar to CetZ proteins. It is also speculated that the possible cell enlargement and membrane complexity during Eukaryogenesis may have necessitated the emergence of a more rigid filamentous structure and led to the evolution of these tubulin-like proteins^43^. Future structural and biochemical studies on synthetic chimeras of Odin FtsZs and OdinTubulin should shed light on the evolution and stabilization of straight protofilaments of tubulin/FtsZ into tubules. An understanding of the cellular functions of these proteins is majorly hindered due to the lack of available culturing techniques for Asgard archaea. However, recent studies on the isolation of members of Lokiarchaeota^3,67^ have opened new avenues for understanding the cellular roles of these Asgard proteins. Thus, future developments in culturing methods, advanced sequencing methods and genetic tools will help build a better understanding of the cell biology of the Asgard archaeal phylum with respect to the role of the multiple FtsZs.

## Acknowledgements

We thank the EM facility at the Division of Biological Sciences, Indian Institute of Science. Electron microscopy facilities at the Department of Physics, IISER Pune, Jyotsna Singh and Radha Chouhan, NCCS Pune for SEC MALS facility. A.D. and J.K. acknowledge the fellowship from IISc. AK acknowledges the PhD fellowship from the Department of Biotechnology. SR acknowledges fellowship from DAE. S.D. acknowledge the CSIR for the SRF fellowship. We thank Prof. Buzz Baum, Dr. Alex Bisson and Prof. PN Rangarajan for their feedback. Schematic diagram is created using BioRender. This paper was typeset with the bioRxiv word template by @Chrelli: www.github.com/chrelli/bioRxiv-word-template.

## Author contributions

Research conceived by SP. Conceptualized by SP, PG, RS, and VK. Methodology designed by JK, AU, SD, SP, PG, and VK. JK cloned all the constructs. JK and AU purified the proteins. SD helped in the standardization of protein purification. JK and AU performed experiments and curated the data. JK, AU, SD, and AK acquired negative EM data. AK, SB, and VK acquired cryo-EM data. AK, SB, and VK performed cryo image analysis. VS and SRP conducted phylogenetic analysis. AD and SR provided resources. JK, AU, SD, SP, PG, and RS analyzed the data. SP, PG, and RS validated the findings. JK, SP drafted the original manuscript. AU, AD, PG, RS, VK, and SRP reviewed and edited the manuscript with the help of all other authors. SP, PG, RS, VK, SRP secured funding.

## Materials and Methods

### Phylogenetic analysis

The maximum likelihood tree was generated using the FtsZ protein sequences from 25 Asgard archaea and 2 Euryarchaea (*Methanobacterium alkalithermotolerans* and *Haloferax volcanii*) as outgroups. For SepF phylogenetic tree 22 SepF sequences were selected. These protein sequences were retrieved from the NCBI database using protein BLAST with Odinarchaeota proteins as query sequences. Sequences were aligned with MEGA7 by using MUSCLE alignment^69^. The phylogenetic tree was generated using IQTREE with LG+G4 as the best-fit model^70^. The support values for each clade, shown at the node, are inferred from ML IQ-TREE 10000 ultrafast bootstrap values.

### Plasmid construction

DNA fragments were synthesized by Twist Biosciences. DNA fragments were amplified using PCR and gel extracted. Fragments and cut vectors were assembled using Hi-fi DNA assembly master mix (NEB) (NewEngland BioLabs; cat.no: E2621L). The reaction mixture was transformed into DH5-alpha cells, and plasmids were isolated, confirmed and sequenced.

### Protein expression and purification

Proteins were expressed in BL21(DE3) cells grown in LB media containing 100 µg/ml of ampicillin at OD600 0.6 with 0.5 mM IPTG (isopropyl β-D-1-thiogalactopyranoside) for 4 hours at 30^°^ C post-induction. Cells were harvested by centrifugation for 15 minutes at 5,000 xg at 4^°^ C. Cells were lysed (Lysis Buffer-50 mM Tris pH 8, 200 mM NaCl, 10% glycerol) followed by clarification by centrifugation at 22,000 xg at 4^°^C. The supernatant was passed through a 5 ml His-Trap column (Cytiva HisTrap HP; 5 ml) pre-equilibrated with a binding buffer (50 mM Tris pH 8, 200 mM NaCl, 10 mM imidazole). Protein was eluted with increasing concentration of imidazole. Elution fractions were loaded onto 12% SDS-PAGE gel, and pure fractions were pooled. Fractions were subjected to cleavage with Ulp protease (1:100 moles ratio of protease: Protein) for 2 hrs (overnight cleavage for OdinFtsZ1). The protein buffer was then exchanged with an imidazole-free buffer (50 mM Tris pH 8, 200 mM NaCl). This was then loaded back to the Ni-NTA column pre-equilibrated with the same buffer to remove the cleaved SUMO tag and the un-cleaved protein. Untagged pure protein was collected in flow through, and then the buffer was exchanged with storage buffer (50 mM HEPES pH 7.4, 50 mM KCl). Protein was concentrated using centricon (10 kDa), flash-frozen and stored at -80^°^C. Cytiva Mono Q column (Mono Q™ 4.6/100 PE, Mono Q™ 10/100 GL) was used for ion exchange chromatography. 50 mM HEPES pH 7.4, 50 mM KCl served as the binding buffer for MonoQ and protein was eluted using a linear gradient of 0 to 40 % of elution buffer (50 mM HEPES pH 7.4, 1 M KCl). After ion exchange chromatography, required fractions were pooled and dialyzed against the storage buffer with a final KCl concentration of 50 mM.

### Light scattering assay

To monitor the polymerization of OdinFtsZ1 and OdinFtsZ2, 90^°^ angle light scattering (Horiba Fluoromax-4) was used. The 200 µL reaction mix contained the buffer (50 mM HEPES, pH 7.4, 50 mM KCl), 2 mM GTP, 5 mM MgCl_2_ and protein at various concentrations. The emission and excitation wavelength for the assay was set to 350 nm, and the slit width used was 1.5 nm. The reaction mix (without GTP) was added to a cuvette with a 1 cm path length and was placed into a temperature-controlled cuvette chamber set at 30^°^ C. Prior to GTP addition, reading was taken for setting a baseline. After 300 seconds, GTP was added to the cuvette to achieve a final concentration of 2 mM and readings were recorded for 40 minutes. The relative intensity profile is obtained by normalizing all the individual intensity values by the average baseline intensity obtained by averaging intensities from 1-300 s.

### Pelleting assay

Purified proteins OdinFtsZ1 and OdinFtsZ2 were spun at 22,000 xg for 20 minutes at 4^°^C initially to remove any aggregates. The protein concentration of the supernatant was then estimated using the Bradford assay^71^. The reaction volume for the pelleting assay was 50 µL, which consisted of 10 µM of protein, 3 mM GTP, 5 mM MgCl_2_ and buffer (50 mM HEPES, pH 7.4, 50 mM KCl). Pelleting of the proteins was monitored in both absence and presence of GTP. Also, both proteins were added in equimolar concentration (10 µM) to monitor co-pelleting. Once the reactions were prepared in the ultracentrifuge (Beckman Coulter Optima MAX-XP Ultracentrifuge) tubes, they were kept inside the rotor (TLA 120.2) set at 30^°^ C for 5 minutes and followed by centrifugation at 100,000 xg for 15 minutes. After the spin, the supernatant was transferred to a fresh vial and the pellet was washed gently with 100 µL buffer (50 mM HEPES pH 7.4, 50 mM KCl). Then, the pellet was resuspended in 50 µL of 5x SDS-PAGE running buffer (125 mM Tris base, 962 mM Glycine and 1.73 mM SDS). Equal volumes of supernatant and pellet fractions were mixed with 2x Laemmli buffer and loaded onto a 12 % SDS-PAGE gel to visualize the proteins. Protein band intensities were analyzed using Fiji (ImageJ2) analysis software^72^. The intensity of bands was calculated as the difference in intensity of a band from that of the average intensity of the background. The percentage of protein in the pellet fraction was calculated by dividing the intensity of the pellet fraction by the sum of the intensities of the pellet and supernatant fractions.

### Pull-down assay for FtsZ

To test the interaction between 6HIS-SUMO tagged OdinFtsZ with OdinSepF, purified OdinFtsZ1 and OdinFtsZ2 were bound to Ni-NTA resin and incubated with untagged OdinSepF. The total reaction volume was 500 µL of binding buffer (50 mM Tris pH 8, 200 mM NaCl) with equimolar amounts of both the proteins, incubated at 4^°^C for 1 hour. After incubation, the beads were washed (thrice) with a wash buffer (50 mM Tris pH 8, 200 mM NaCl, 25 mM imidazole) to remove non-specific binding. Proteins bound with Ni-NTA beads were then eluted with a 500 µL elution buffer (50 mM Tris pH 8, 200 mM NaCl, 250 mM Imidazole). A control experiment with 6HIS-SUMO was done to eliminate the possible SUMO-based binding. Each fraction was collected and loaded on 12% SDS-PAGE gel to visualize the proteins.

### Liposome preparation

Lipids utilized in all the liposome-based experiments were acquired from Avanti Polar Lipids, namely, dioleoyl phosphatidylglycerol (DOPG; 840475C) and dioleoyl phosphatidylcholine (DOPC; 850375C). To make a 2 mM stock solution of lipids, the required amount of lipid in chloroform was aliquoted in a small test tube and then dried with nitrogen (or argon) to remove chloroform. After this, the dried lipids were resuspended in HK50 buffer (50 mM HEPES pH 7.4, 50 mM KCl), such that the final stock concentration was 2 mM. This lipid solution was then extruded through a 100 nm polycarbonate membrane (Avanti Polar Lipids) and used for all the liposome-based assays.

### Liposome pelleting assay

Purified proteins were spun at 100,000 xg for 15 minutes at 4^°^C to remove any aggregates. The concentration of supernatant was then estimated using the Bradford assay. The reaction volume used for the assay was 100 µL, which consisted of 2 µM protein, and varying concentrations of liposome (0 - 600 µM) in HK50 buffer. 100 µL reaction was made in an ultracentrifuge tube and incubated for 15 minutes at 30^°^C. Then, the reactions were spun at 100,000 xg for 15 minutes. After the spin, the supernatant was transferred to a fresh vial and the pellet was washed gently with 200 µL HK50 buffer and resuspended in 50 µl of the same buffer. Equal volumes of supernatant and pellet fractions were mixed with 2x Laemmli buffer and loaded onto a 12 % SDS-PAGE gel to visualize the proteins. Protein band intensities were analyzed using Fiji (ImageJ2)^72^. The intensity of bands was calculated as the difference in intensity of a band from that of the average intensity of the background. The percentage of protein in the pellet fraction was calculated by dividing the intensity of the pellet fraction by the sum of the intensities of the pellet and supernatant fractions, and loaded on 15% SDS-PAGE gel to visualize the proteins.

### Structural analysis

All the AlphaFold-predicted structures, *E. coli* FtsZ crystal structures (PDB ID: 6UMK) and HvCetZ1 (PDB: 4B46) were downloaded from UniProt for structural analysis. Superimposition of structures and the rest of the structural analysis were done using ChimeraX^73^. Web logo for sequence conservation was made using a web logo (https://weblogo.berkeley.edu/logo.cgi). HeliQuest^74^ was used for amphipathic helix prediction, helical wheel representation and calculation of hydrophobic moment and hydrophobicity parameters.

### Size Exclusion Chromatography coupled with Multi-Angle Light Scattering (SEC-MALS)

SEC-MALS experiment enabled the accurate mass estimation for purified OdinSepF. Superdex 75 Increase 10/300 GL column was used for SEC, which was connected to an Agilent HPLC unit with an 18-angle light scattering detector (Wyatt Dawn HELIOS II) and a refractive index detector (Wyatt Optilab T-rEX). The experiments were performed at room temperature. The column was equilibrated with HK50 buffer at a flow rate of 0.4 ml/min. BSA at 2 mg/ml was used to calibrate the system. Purified OdinSepF (2 mg/ml, 100 µL) was subsequently loaded to estimate the molecular weight of the eluted peaks. The Zimm model implemented in ASTRA software^75^ was used for the curve fitting and estimation of molecular weights. GraphPad Prism was used to plot the average molar mass from fitted plots.

### Negative staining

The protein samples were thawed, diluted in 50 mM HEPES pH 7.4, 50 mM KCl, 5mM MgCl_2_ and incubated with a final concentration of 2 mM nucleotide (GTP, GTPγS, GDP or GMPCPP) at 30°C for 15 minutes or as indicated. The final concentration of OdinFtsZ1 was 0.04 mg/ml, and that of OdinFtsZ2 was 0.1 mg/ml for all conditions. Maxtaform Cu/Rh 3 mm grids were coated with homemade carbon film on one side. The carbon-coated side was glow discharged in a Quorum GloQube glow discharge instrument at 20 mA for 10 seconds. 3 µL of the protein sample was applied to the grid, incubated for a minute, and washed three times with MilliQ water. The excess water was blotted with Whatman filter paper No. 1 and stained with 2% uranyl acetate twice with a final incubation for 1 minute, after which the excess stain was removed and air dried for 1 minute. The grids were then imaged using Talos F200S G2 TEM operated at 200 kV and a CETA camera.

### Cryo-EM Sample preparation, data collection and image processing

OdinFtsZ1 and OdinFtsZ2 with 2 mM GTP were prepared similar to negative staining protocol and applied to glow-discharged Quantifoil R 0.6/1 (Au, 300 mesh) grids held in a Vitrobot Mark IV (ThermoFisher Scientific) at 20°C and 100 % relative humidity. Grids were blotted for 3-4 seconds and rapidly frozen in liquid ethane and transferred to liquid nitrogen until imaged in the microscope. For imaging, grids were transferred to Krios cartridges and loaded onto Titan Krios TEM (ThermoFisher Scientific). Grids were screened for good ice thickness and particle distribution and data was collected at a nominal magnification of 75,000 x corresponding to a pixel size of 1.07 Å with a Falcon 3 detector. Image processing was performed with RELION and CryoSPARC^76–78^. The alignment of the movie frames was performed with the MotionCorr routine in RELION, followed by CTF estimation with CTFFIND4^79^. For OdinFtsZ2, initial particle picking was performed manually and the classes obtained were used for template-based picking. For OdinFtsZ1, filament tracer routing in CryoSPARC was used. The 2D classification was performed using default parameters in RELION and CryoSPARC.

---

**Figure S1:**
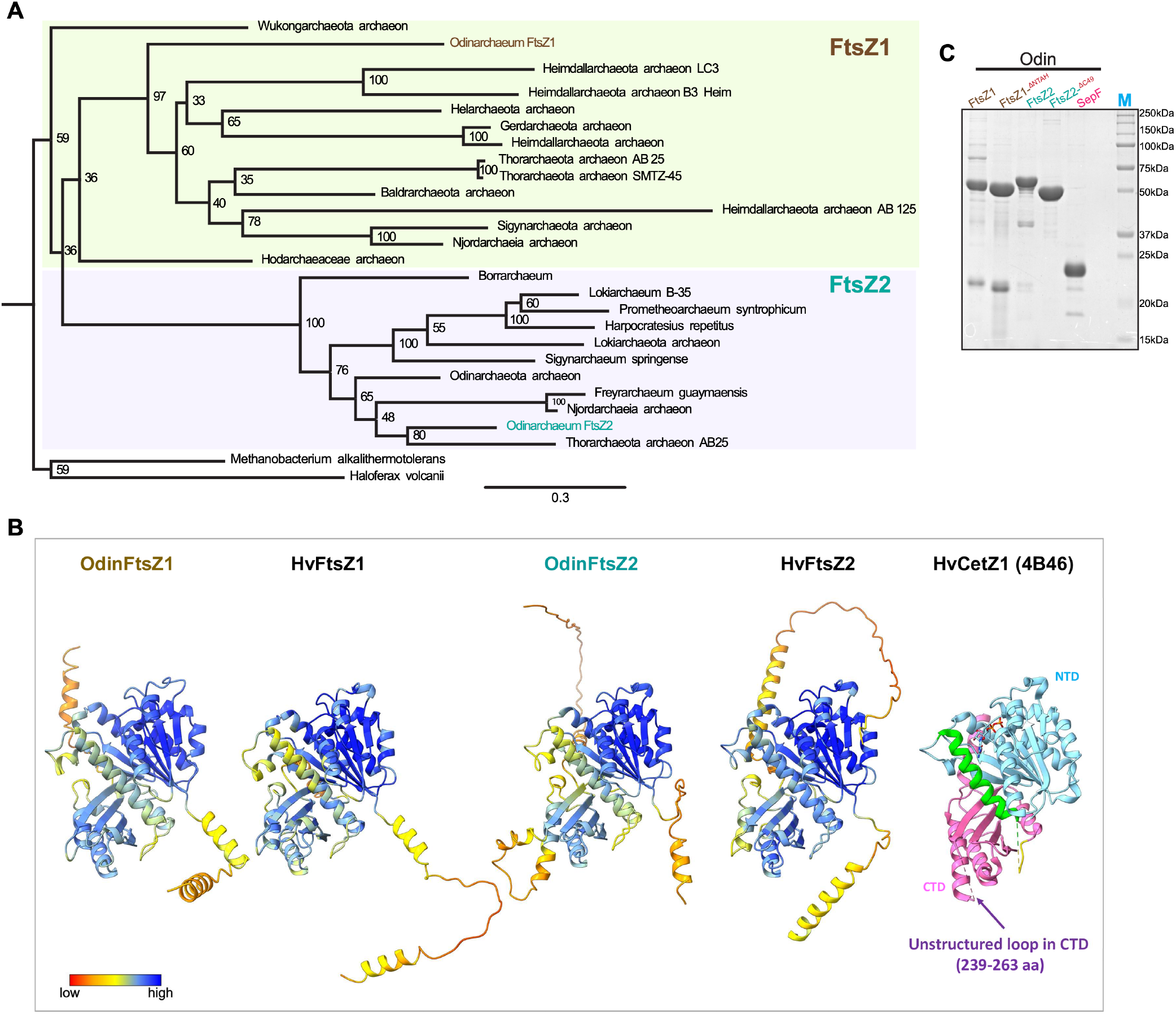
Sequence and structural features and purification profile of OdinFtsZs. **(A)** Maximum likelihood tree for 25 representative FtsZ sequences of Asgard archaeon along with *Methanobacterium alkalithermotolerans* and *Haloferax volcanii* as outgroups. **(B)** Protomer structure comparison of OdinFtsZ1 and OdinFtsZ2 monomers with *Haloferax volcanii* (HvFtsZ1 HvFtsZ2, and HvCetZ). The cartoon representation of the structures is coloured according to the pLDDT score from AlphaFold2. **(C)** Representative 12% SDS-PAGE gel for all the untagged proteins used in the study.

**Figure S2:**
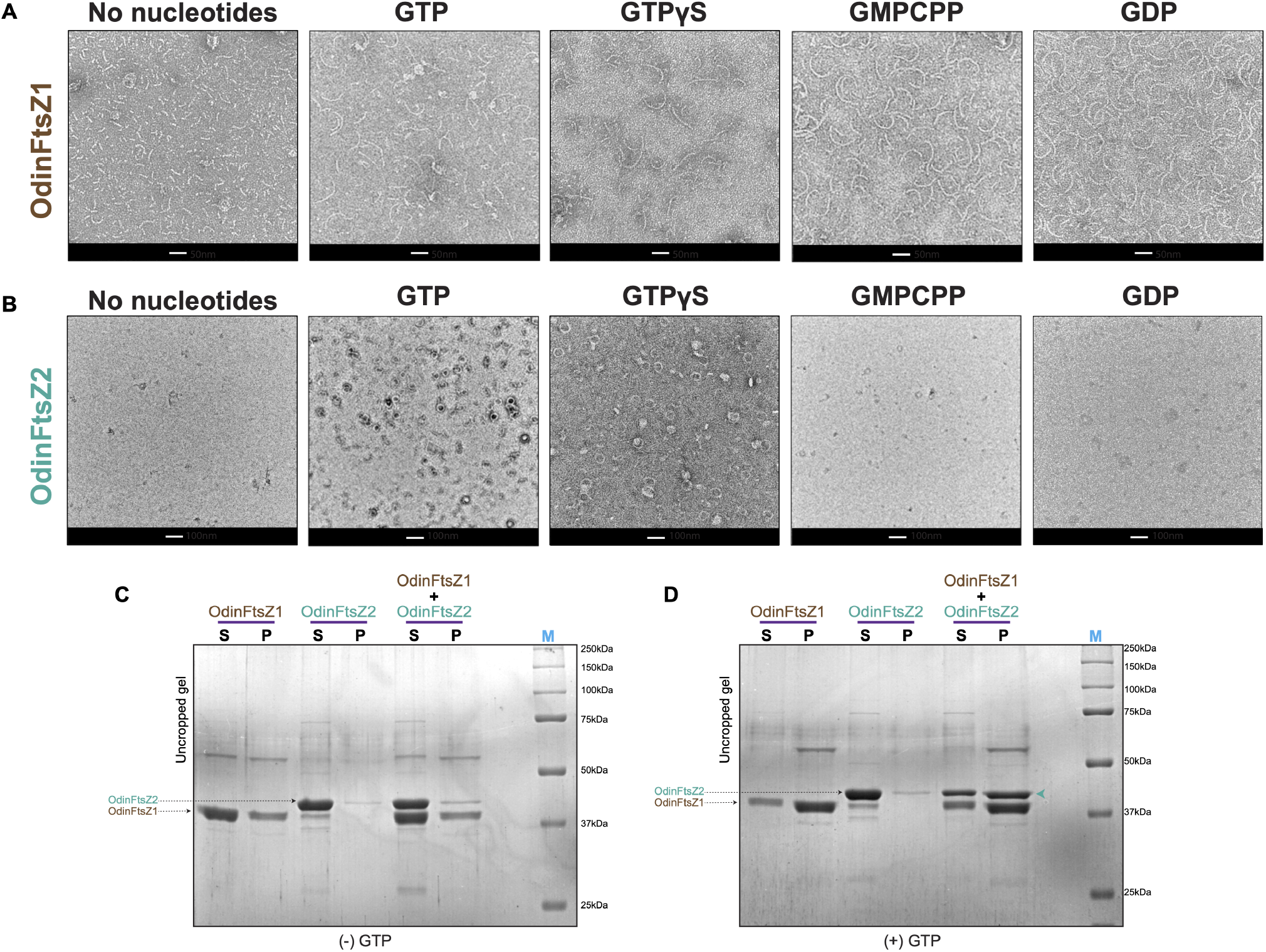
Filament assembly of OdinFtsZs. **(A)** Negative stain micrographs of 0.04 mg ml^−1^ OdinFtsZ1 supplemented with (a) no nucleotide, (b) 2 mM GTP, (c) 2 mM GTPγS, (d) 2 mM GMPCPP and (e) 2 mM GDP. The scale bar represents 50 nm. **(B)** Negative stain micrographs of 0.04 mg ml^−1^ OdinFtsZ2 supplemented with (a) no nucleotide, (b) 2 mM GTP, (c) 2mM GTPγS, (d) 2mM GMPCPP and (e) 2 mM GDP. The scale bar represents 100 nm. Each experiment was repeated independently at least twice. **(C)** Uncropped Coomassie Brilliant Blue (CBB) gel of Fig. 2E (top) without the addition of GTP. **(D)** Uncropped CBB gel of Fig. 2E (bottom) with the addition of 5 mM GTP.

**Figure S3:**
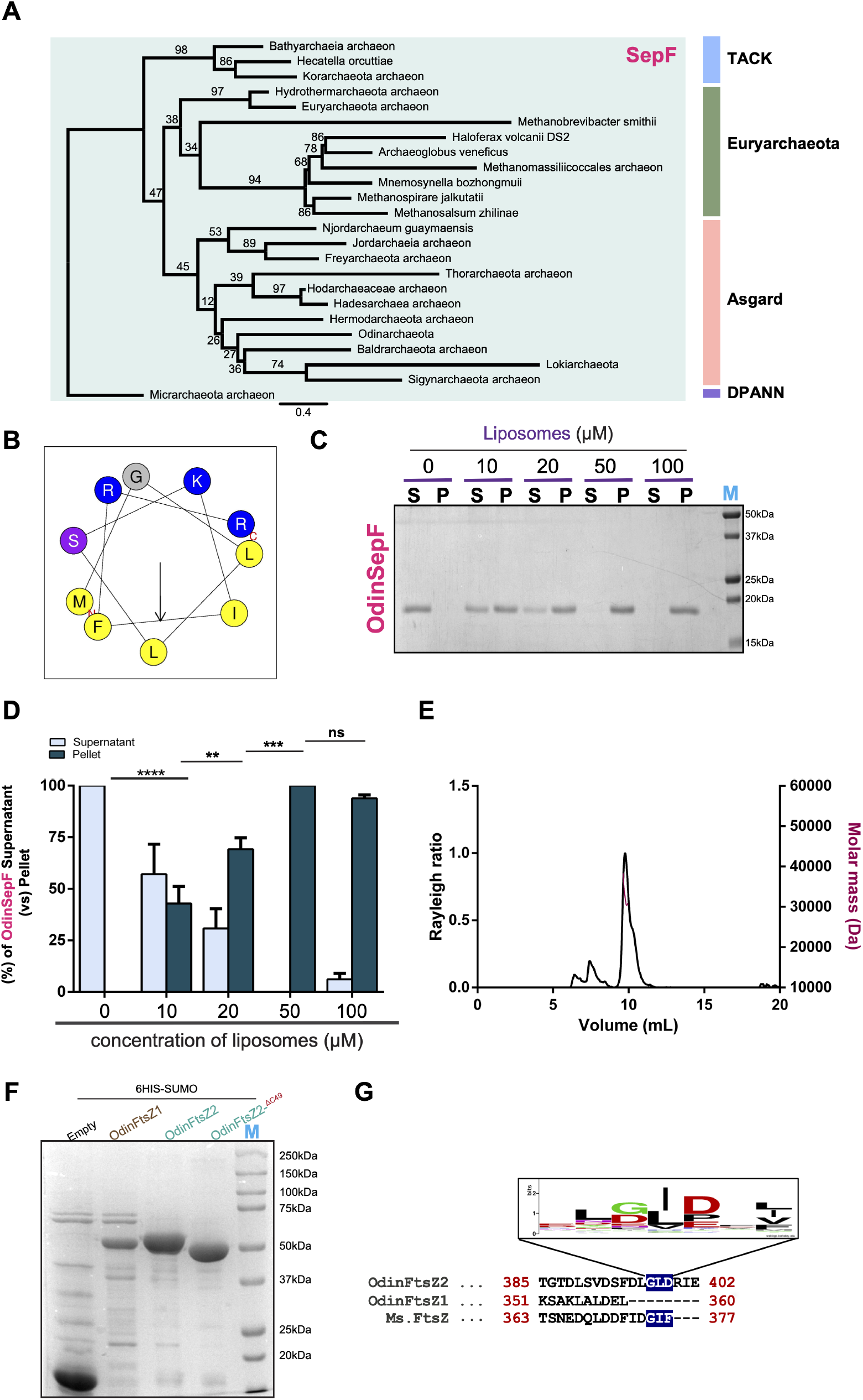
Characteristic features of OdinSepF. **(A)** Maximum likelihood tree for 22 representatives archaeal SepF sequences. **(B)** Representative helical wheel diagram for the predicted AH at the N-terminus of OdinSepF screened using Heliquest (version 1.2 Analysis module)^68^. **(C)** Representative 15% SDS-PAGE gel for OdinSepF sedimentation with increasing liposome concentration. The supernatant (S) and pellet (P) fractions were analyzed and loaded upon sedimentation at 100,000 ×*g*. Lanes 1 and 2 represent S and P for only OdinSepF without liposomes. **(D)** Representative plot for the percentage fraction of OdinSepF in the supernatant (S) and pellet (P) fractions with increasing concentrations of liposomes as represented in Fig. S3C. The pellet (or supernatant) fraction intensities corresponding to the OdinFtsZ1 band were calculated as the intensity of the pellet (or supernatant) divided by the sum of the two intensities and represented in the graph as a percentage. (N=3, Two-Way ANOVA with Sidak’s multiple comparisons test was used, ^ns^ non-significant, ** p - 0.0055, *** p < 0.001, **** p < 0.0001) **(E)** SEC-MALS profile of purified OdinSepF. The chromatogram displays the calculated molar mass of the peaks (Da) and Rayleigh ratio as red and black lines, respectively. **(F)** Representative 15% SDS-PAGE gel for proteins used in Fig. 3A, 3B **(G)** Sequence logo for the conservation of the GID motif in the C-terminal tail of FtsZ^14^.

**Supplementary Table 1.** List of plasmids used in this study.

| **Plasmid Number** | **Description** | **Source** |
| --- | --- | --- |
| piSP65 | pET28a-6HIS-bdSUMO (Empty) | Hatano et.al 2022<sup>8</sup> |
| piSP1367 | pET28a-6HIS-bdSUMO-OdinFtsZ1 | This Study |
| piSP1368 | pET28a-6HIS-bdSUMO-OdinFtsZ2 | This Study |
| piSP1753 | pET28a-6HIS-bdSUMO-OdinSepF | This Study |
| piSP1780 | pET28a-6HIS-bdSUMO-OdinFtsZ1(ΔNTAH) (14-363 a.a) | This Study |
| piSP1786 | pET28a-6HIS-bdSUMO-OdinFtsZ2<sup>ΔC49</sup>(1-355 a.a) | This Study |

## References

1. Zaremba-Niedzwiedzka, K. et al. Asgard archaea illuminate the origin of eukaryotic cellular complexity. Nature 541, 353–358 (2017).

2. Spang, A. et al. Complex archaea that bridge the gap between prokaryotes and eukaryotes. Nature 521, 173–179 (2015).

3. Imachi, H. et al. Isolation of an archaeon at the prokaryote–eukaryote interface. Nature 577, 519–525 (2020).

4. MacLeod, F. et al. Asgard archaea: Diversity, function, and evolutionary implications in a range of microbiomes. AIMS Microbiol. 5, 48–61 (2019).

5. Eme, L. et al. Inference and reconstruction of the heimdallarchaeial ancestry of eukaryotes. Nature 618, 992–999 (2023).

6. Akıl, C. & Robinson, R. C. Genomes of Asgard archaea encode profilins that regulate actin. Nature 562, 439–443 (2018).

7. Akıl, C. et al. Insights into the evolution of regulated actin dynamics via characterization of primitive gelsolin/cofilin proteins from Asgard archaea. Proc. Natl. Acad. Sci. 117, 19904–19913 (2020).

8. Hatano, T. et al. Asgard archaea shed light on the evolutionary origins of the eukaryotic ubiquitin-ESCRT machinery. Nat. Commun. 13, 3398 (2022).

9. Melnikov, N. et al. The Asgard archaeal ESCRT-III system forms helical filaments and remodels eukaryotic membranes, shedding light on the emergence of eukaryogenesis. 2022.09.07.506706 Preprint at 10.1101/2022.09.07.506706 (2024).

10. Souza, D. P. et al. Evolutionarily conserved principles of ESCRT-III-mediated membrane remodelling revealed by a two-subunit Asgard archaeal system. 2024.07.01.601590 Preprint at 10.1101/2024.07.01.601590 (2024).

11. Bi, E. & Lutkenhaus, J. FtsZ ring structure associated with division in Escherichia coli. Nature 354, 161–164 (1991).

12. Erickson, H. P., Anderson, D. E. & Osawa, M. FtsZ in Bacterial Cyto-kinesis: Cytoskeleton and Force Generator All in One. Microbiol. Mol. Biol. Rev. 74, 504–528 (2010).

13. Margolin, W. FtsZ and the division of prokaryotic cells and organelles. Nat. Rev. Mol. Cell Biol. 6, 862–871 (2005).

14. Liao, Y., Ithurbide, S., Evenhuis, C., Löwe, J. & Duggin, I. G. Cell division in the archaeon Haloferax volcanii relies on two FtsZ proteins with distinct functions in division ring assembly and constriction. Nat. Microbiol. 6, 594–605 (2021).

15. Li, Z., Trimble, M. J., Brun, Y. V. & Jensen, G. J. The structure of FtsZ filaments in vivo suggests a force-generating role in cell division. EMBO J. 26, 4694–4708 (2007).

16. Bisson-Filho, A. W. et al. Treadmilling by FtsZ filaments drives peptidoglycan synthesis and bacterial cell division. Science 355, 739–743 (2017).

17. Margolis, R. L. & Wilson, L. Microtubule treadmills—possible molecular machinery. Nature 293, 705–711 (1981).

18. McCausland, J. W. et al. Treadmilling FtsZ polymers drive the directional movement of sPG-synthesis enzymes via a Brownian ratchet mechanism. Nat. Commun. 12, 609 (2021).

19. Whitley, K. D. et al. FtsZ treadmilling is essential for Z-ring condensation and septal constriction initiation in Bacillus subtilis cell division. Nat. Commun. 12, 2448 (2021).

20. McQuillen, R. & Xiao, J. Insights into the Structure, Function, and Dy-namics of the Bacterial Cytokinetic FtsZ-Ring. Annu. Rev. Biophys. 49, 309–341 (2020).

21. Mukherjee, A. & Lutkenhaus, J. Dynamic assembly of FtsZ regulated by GTP hydrolysis. EMBO J. 17, 462–469 (1998).

22. Oliva, M. A. et al. Assembly of archaeal cell division protein FtsZ and a GTPase-inactive mutant into double-stranded filaments. J. Biol. Chem. 278, 33562–33570 (2003).

23. Lu, C., Reedy, M. & Erickson, H. P. Straight and Curved Conformations of FtsZ Are Regulated by GTP Hydrolysis. J. Bacteriol. 182, 164–170 (2000).

24. Chakraborty, J. et al. Dynamics of interdomain rotation facilitates FtsZ filament assembly. J. Biol. Chem. 300, (2024).

25. Housman, M., Milam, S. L., Moore, D. A., Osawa, M. & Erickson, H. P. FtsZ Protofilament Curvature Is the Opposite of Tubulin Rings. Biochemistry 55, 4085 (2016).

26. Du, S., Pichoff, S., Kruse, K. & Lutkenhaus, J. FtsZ filaments have the opposite kinetic polarity of microtubules. Proc. Natl. Acad. Sci. 115, 10768–10773 (2018).

27. Osawa, M. & Erickson, H. P. Liposome division by a simple bacterial division machinery. Proc. Natl. Acad. Sci. 110, 11000–11004 (2013).

28. Szwedziak, P., Wang, Q., Bharat, T. A. M., Tsim, M. & Löwe, J. Architecture of the ring formed by the tubulin homologue FtsZ in bacterial cell division. eLife 3, e04601 (2014).

29. Osawa, M., Anderson, D. E. & Erickson, H. P. Reconstitution of Contractile FtsZ Rings in Liposomes. Science 320, 792–794 (2008).

30. Bork, P., Sander, C. & Valencia, A. An ATPase domain common to prokaryotic cell cycle proteins, sugar kinases, actin, and hsp70 heat shock proteins. Proc. Natl. Acad. Sci. 89, 7290–7294 (1992).

31. Pichoff, S. & Lutkenhaus, J. Tethering the Z ring to the membrane through a conserved membrane targeting sequence in FtsA. Mol. Microbiol. 55, 1722–1734 (2005).

32. Szwedziak, P., Wang, Q., Freund, S. M. & Löwe, J. FtsA forms actinlike protofilaments. EMBO J. 31, 2249–2260 (2012).

33. Hale, C. A. & de Boer, P. A. Direct binding of FtsZ to ZipA, an essential component of the septal ring structure that mediates cell division in E. coli. Cell 88, 175–185 (1997).

34. Hamoen, L. W., Meile, J.-C., De Jong, W., Noirot, P. & Errington, J. SepF, a novel FtsZ-interacting protein required for a late step in cell division. Mol. Microbiol. 59, 989–999 (2006).

35. Duman, R. et al. Structural and genetic analyses reveal the protein SepF as a new membrane anchor for the Z ring. Proc. Natl. Acad. Sci. 110, E4601–E4610 (2013).

36. Pende, N. et al. SepF is the FtsZ anchor in archaea, with features of an ancestral cell division system. Nat. Commun. 12, 3214 (2021).

37. Nußbaum, P., Gerstner, M., Dingethal, M., Erb, C. & Albers, S.-V. The archaeal protein SepF is essential for cell division in Haloferax volcanii. Nat. Commun. 12, 3469 (2021).

38. Nußbaum, P. et al. Proteins containing photosynthetic reaction centre domains modulate FtsZ-based archaeal cell division. Nat. Microbiol. 9, 698–711 (2024).

39. Lindås, A.-C., Karlsson, E. A., Lindgren, M. T., Ettema, T. J. G. & Bernander, R. A unique cell division machinery in the Archaea. Proc. Natl. Acad. Sci. U. S. A. 105, 18942–18946 (2008).

40. Samson, R. Y., Obita, T., Freund, S. M., Williams, R. L. & Bell, S. D. A Role for the ESCRT System in Cell Division in Archaea. Science 322, 1710–1713 (2008).

41. Hurtig, F. et al. The patterned assembly and stepwise Vps4-mediated disassembly of composite ESCRT-III polymers drives archaeal cell division. Sci. Adv. 9, eade5224 (2023).

42. Tamarit, D. et al. A closed Candidatus Odinarchaeum chromosome exposes Asgard archaeal viruses. Nat. Microbiol. 7, 948–952 (2022).

43. Akıl, C. et al. Structure and dynamics of Odinarchaeota tubulin and the implications for eukaryotic microtubule evolution. Sci. Adv. 8, eabm2225 (2022).

44. Larsen, R. A. et al. Treadmilling of a prokaryotic tubulin-like protein, TubZ, required for plasmid stability in Bacillus thuringiensis. Genes Dev. 21, 1340 (2007).

45. Chaikeeratisak, V. et al. The Phage Nucleus and Tubulin Spindle Are Conserved among Large Pseudomonas Phages. Cell Rep. 20, 1563– 1571 (2017).

46. Erb, M. L. et al. A bacteriophage tubulin harnesses dynamic instability to center DNA in infected cells. eLife 3, e03197 (2014).

47. Duggin, I. G. et al. CetZ tubulin-like proteins control archaeal cell shape. Nature 519, 362–365 (2015).

48. Nußbaum, P. et al. An Oscillating MinD Protein Determines the Cellular Positioning of the Motility Machinery in Archaea. Curr. Biol. 30, 4956-4972.e4 (2020).

49. Jumper, J. et al. Highly accurate protein structure prediction with AlphaFold. Nature 596, 583–589 (2021).

50. Vaughan, S., Wickstead, B., Gull, K. & Addinall, S. G. Molecular Evolution of FtsZ Protein Sequences Encoded Within the Genomes of Archaea, Bacteria, and Eukaryota. J. Mol. Evol. 58, 19–29 (2004).

51. The essential bacterial cell-division protein FtsZ is a GTPase | Nature. https://www.nature.com/articles/359254a0.

52. RayChaudhuri, D. & Park, J. T. Escherichia coli cell-division gene ftsZ encodes a novel GTP-binding protein. Nature 359, 251–254 (1992).

53. Erickson, H. P., Taylor, D. W., Taylor, K. A. & Bramhill, D. Bacterial cell division protein FtsZ assembles into protofilament sheets and minirings, structural homologs of tubulin polymers. Proc. Natl. Acad. Sci. 93, 519–523 (1996).

54. Mukherjee, A. & Lutkenhaus, J. Analysis of FtsZ Assembly by Light Scattering and Determination of the Role of Divalent Metal Cations. J. Bacteriol. 181, 823–832 (1999).

55. Romberg, L., Simon, M. & Erickson, H. P. Polymerization of FtsZ, a Bacterial Homolog of Tubulin: IS ASSEMBLY COOPERATIVE?*. J. Biol. Chem. 276, 11743–11753 (2001).

56. González, J. M. et al. Cooperative behavior of Escherichia coli cell-division protein FtsZ assembly involves the preferential cyclization of long single-stranded fibrils. Proc. Natl. Acad. Sci. 102, 1895–1900 (2005).

57. Wagstaff, J. M. et al. A Polymerization-Associated Structural Switch in FtsZ That Enables Treadmilling of Model Filaments. mBio 8, 10.1128/mbio.00254-17 (2017).

58. Fujita, J. et al. Structures of a FtsZ single protofilament and a doublehelical tube in complex with a monobody. Nat. Commun. 14, 4073 (2023).

59. Cendrowicz, E. et al. Bacillus subtilis SepF Binds to the C-Terminus of FtsZ. PLOS ONE 7, e43293 (2012).

60. Structural and genetic analyses reveal the protein SepF as a new membrane anchor for the Z ring. https://www.pnas.org/doi/10.1073/pnas.1313978110 doi:10.1073/pnas.1313978110.

61. Sogues, A. et al. Essential dynamic interdependence of FtsZ and SepF for Z-ring and septum formation in Corynebacterium glutamicum. Nat. Commun. 11, 1641 (2020).

62. Dutta, S., Poddar, S., Chakraborty, J., Srinivasan, R. & Gayathri, P. An amphipathic helix facilitates direct membrane binding of Mycoplasma FtsZ. 2023.08.29.555414 Preprint at 10.1101/2023.08.29.555414 (2023).

63. novel amphiphilic motif at the C-terminus of FtsZ1 facilitates chloroplast division | The Plant Cell | Oxford Academic. https://aca-demic.oup.com/plcell/article/34/1/419/6424909.

64. An, J., Wang, L., Hong, C. & Gao, H. Evolution and Functional Differentiation of the C-terminal Motifs of FtsZs During Plant Evolution. Mol. Biol. Evol. 41, msae145 (2024).

65. Erickson, H. P. Evolution of the cytoskeleton. BioEssays News Rev. Mol. Cell. Dev. Biol. 29, 668 (2007).

66. Krupka, M. et al. Escherichia coli FtsA forms lipid-bound minirings that antagonize lateral interactions between FtsZ protofilaments. Nat. Commun. 8, 15957 (2017).

67. Rodrigues-Oliveira, T. et al. Actin cytoskeleton and complex cell architecture in an Asgard archaeon. Nature 613, 332–339 (2023).

68. HeliQuest ComputParam form version3. https://heliquest.ipmc.cnrs.fr/cgi-bin/ComputParams.py.

69. Kumar, S., Stecher, G. & Tamura, K. MEGA7: Molecular Evolutionary Genetics Analysis Version 7.0 for Bigger Datasets. Mol. Biol. Evol. 33, 1870–1874 (2016).

70. Trifinopoulos, J., Nguyen, L.-T., von Haeseler, A. & Minh, B. Q. W-IQ-TREE: a fast online phylogenetic tool for maximum likelihood analysis. Nucleic Acids Res. 44, W232–235 (2016).

71. Bradford, M. M. A rapid and sensitive method for the quantitation of microgram quantities of protein utilizing the principle of protein-dye binding. Anal. Biochem. 72, 248–254 (1976).

72. Schindelin, J. et al. Fiji: an open-source platform for biological-image analysis. Nat. Methods 9, 676–682 (2012).

73. Goddard, T. D. et al. UCSF ChimeraX: Meeting modern challenges in visualization and analysis. Protein Sci. Publ. Protein Soc. 27, 14–25 (2018).

74. HELIQUEST: a web server to screen sequences with specific α-helical properties | Bioinformatics | Oxford Academic. https://academic.oup.com/bioinformatics/article/24/18/2101/192677?login=true.

75. ASTRA. https://www.wyatt.com/products/software/astra.html (2019).

76. Scheres, S. H. W. RELION: Implementation of a Bayesian approach to cryo-EM structure determination. J. Struct. Biol. 180, 519–530 (2012).

77. cryoSPARC: algorithms for rapid unsupervised cryo-EM structure determination | Nature Methods. https://www.nature.com/articles/nmeth.4169.

78. Zivanov, J. et al. New tools for automated high-resolution cryo-EM structure determination in RELION-3. eLife 7, e42166 (2018).

79. Rohou, A. & Grigorieff, N. CTFFIND4: Fast and accurate defocus estimation from electron micrographs. J. Struct. Biol. 192, 216–221 (2015).

