## Supplementary figures and images for "Distinct filament morphology and membrane tethering features of the dual FtsZs in Odinarchaeota"

### Supplemental Figures

FigureS1

Kumari et al

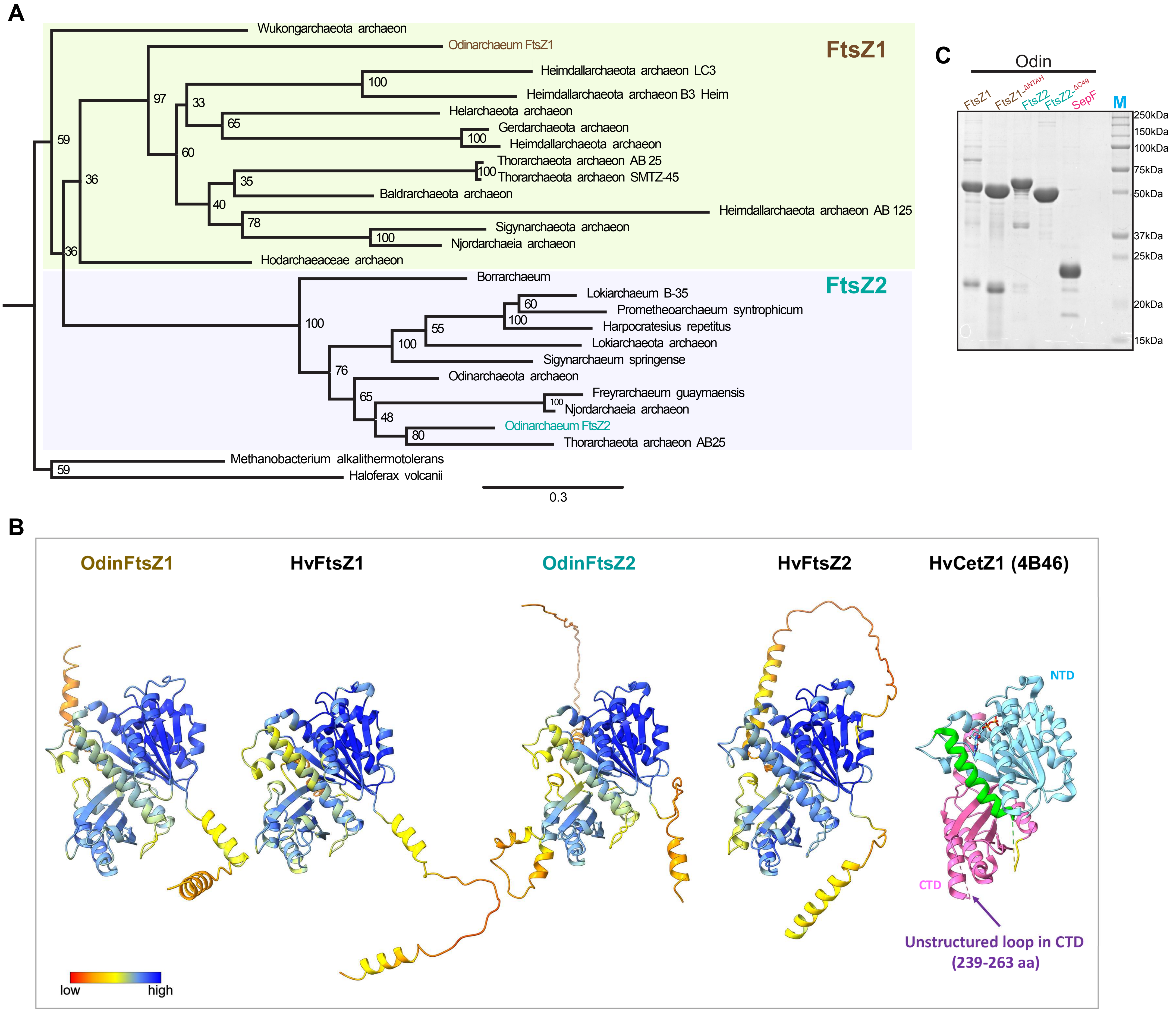

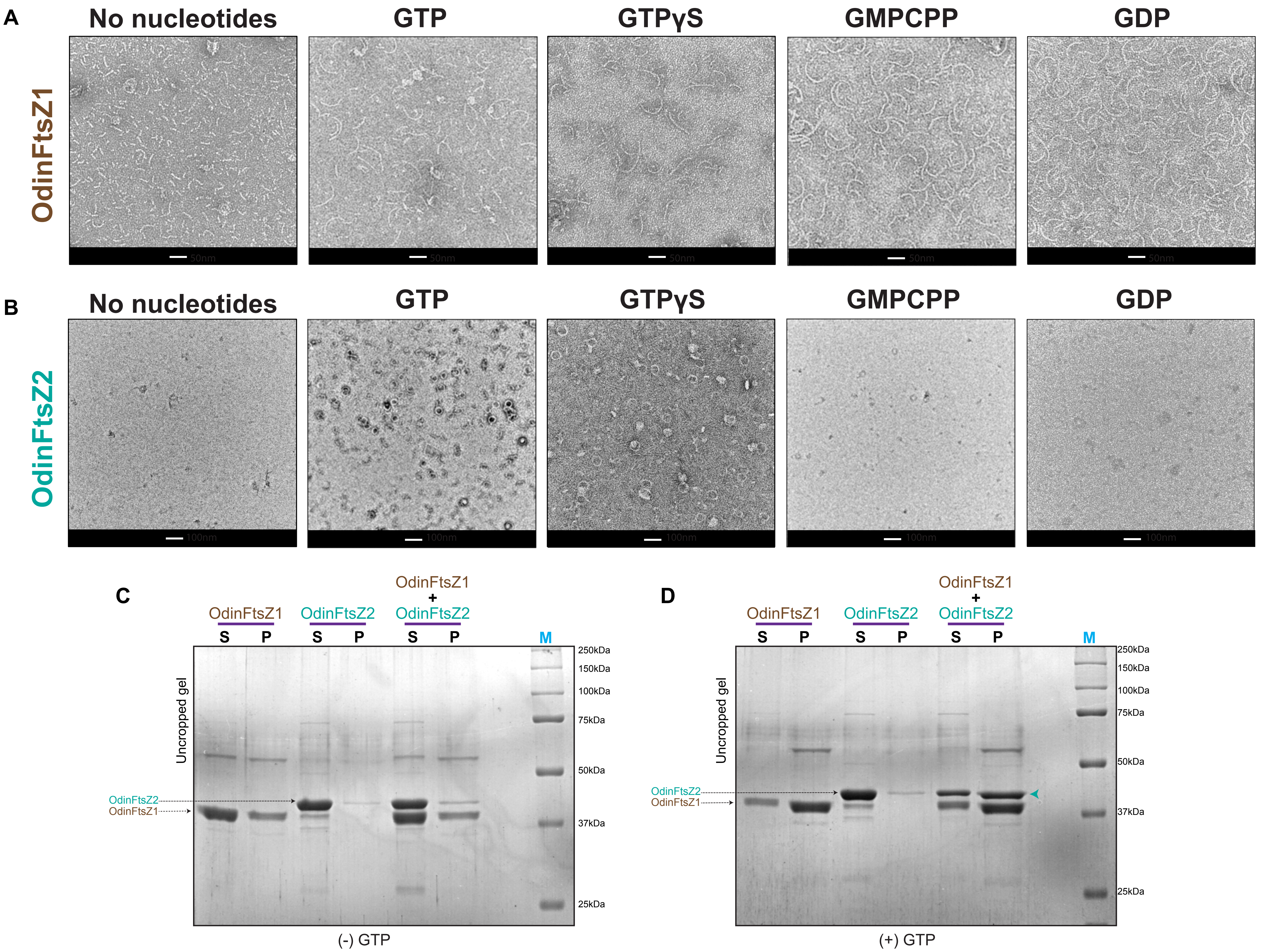

FigureS3

Kumari et al

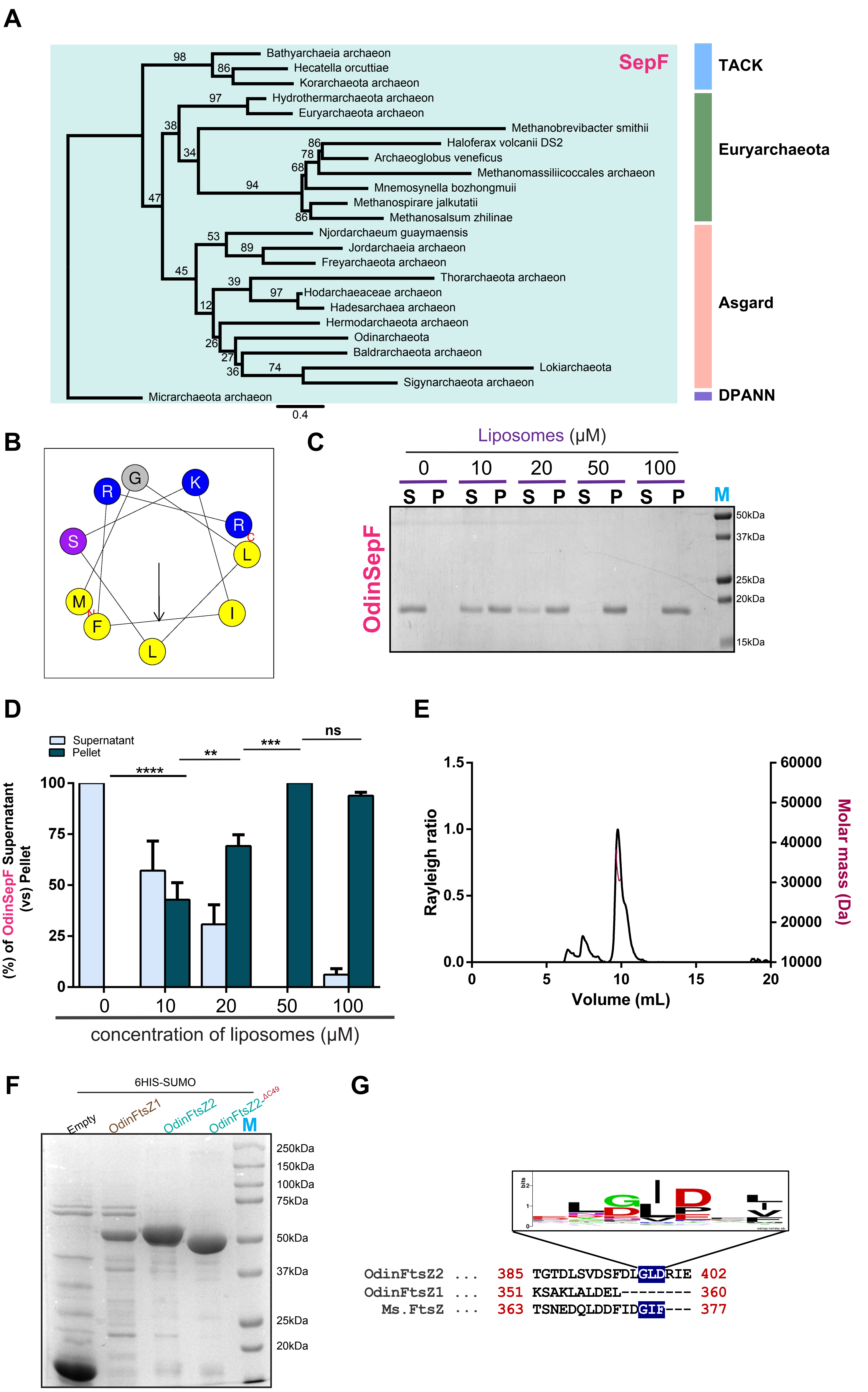
